# Oxidative stress-induced MMP- and γ-secretase-dependent VE-cadherin processing is modulated by the proteasome and BMP9/10

**DOI:** 10.1101/2022.11.23.517709

**Authors:** Caterina Ivaldo, Mario Passalacqua, Anna Lisa Furfaro, Cristina d’Abramo, Santiago Ruiz, Prodyot K. Chatterjee, Christine N. Metz, Mariapaola Nitti, Philippe Marambaud

## Abstract

Classical cadherins, including vascular endothelial (VE)-cadherin, are targeted by matrix metalloproteinases (MMPs) and γ-secretase during adherens junction (AJ) disassembly, a mechanism that might have relevance for endothelial cell (EC) integrity and vascular homeostasis. Here, we show that oxidative stress triggered by H_2_O_2_ exposure induced efficient VE-cadherin proteolysis by MMPs and γ-secretase in human umbilical endothelial cells (HUVECs). The cytoplasmic domain of VE-cadherin produced by γ-secretase, VE-Cad/CTF2 - a fragment that has eluded identification so far - could readily be detected after H_2_O_2_ treatment. VE-Cad/CTF2, released into the cytosol, was tightly regulated by proteasomal degradation and was sequentially produced from an ADAM10/17-generated C-terminal fragment, VE-Cad/CTF1. Interestingly, BMP9 and BMP10, two circulating ligands critically involved in vascular maintenance, significantly reduced VE-Cad/CTF2 levels during H_2_O_2_ challenge, as well as mitigated H_2_O_2_-mediated actin cytoskeleton disassembly during VE-cadherin processing. Notably, BMP9/10 pretreatments efficiently reduced apoptosis induced by H_2_O_2_, favoring endothelial cell recovery. Thus, oxidative stress is a trigger of MMP- and γ-secretase-mediated endoproteolysis of VE-cadherin and AJ disassembly from the cytoskeleton in ECs, a mechanism that is negatively controlled by the EC quiescence factors, BMP9 and BMP10.

## Introduction

The cadherin superfamily is the major component of the adherens junctions (AJs) in epithelial, neural, and endothelial tissues^1–3^. These cell-cell adhesion proteins are crucial for AJ formation and cell-cell contact organization^4,5^. Cadherin cleavage and AJ disassembly accompany apoptotic cell death^6^ and cell responses to intracellular calcium imbalance^7,8^. Like several other transmembrane proteins, cadherins can undergo endoproteolytic cleavage by the MMPs, ADAM10 and ADAM17, as well as γ-secretase^9,10^. Indeed, it has been shown that under apoptosis or calcium influx, MMPs work in concert with γ-secretase to disassemble AJs and control transcriptional responses by cleaving key members of the classical cadherin family, such as epithelial (E)-cadherin, neural (N)-cadherin, and vascular endothelial (VE)-cadherin^11–13^. We reported that upon calcium influx or apoptosis induction, E-cadherin is proteolyzed by MMPs (later identified as ADAM10 and ADAM17^14^) and γ-secretase to release the C-terminal fragments E-Cad/CTF1 and E-Cad/CTF2, respectively^11^. This proteolytic process triggered AJ disassembly, and γ-secretase proteolysis of E-cadherin specifically facilitated α- and β-catenin release from the cytoskeleton. VE-cadherin is predominantly expressed in endothelial cells (ECs) where it regulates endothelial functions through complex dynamics that involve the binding with different cytoskeleton proteins^15^. Indeed, by binding β-catenin, α-catenin, p120 and plakoglobin, VE-cadherin binds actin filaments to regulate endothelial migration, permeability, vascular remodeling and angiogenesis^16,17^. VE-cadherin was shown to be cleaved by MMPs and γ-secretase to increase vascular permeability^12^.

Oxidative stress (OS) is a common mediator of endothelial dysfunction and EC damage in many diseases, such as diabetes and atherosclerosis^18,19^, and plays crucial roles in vascular aging^20^ and endothelial cell apoptosis^21^. Moreover, OS is involved in the modification of endothelial cell junctions^22^. Indeed, OS in endothelial cells increases permeability through the modification of endothelial junctions and cell contraction^22–25^. In addition, OS favors leukocyte transmigration by remodeling cell-cell junctions via VE-cadherin and β-catenin posttranscriptional modifications^26^. However, the role of OS in the proteolytic processing of VE-cadherin has never been investigated.

The circulating factors, BMP9 and BMP10, bind with high affinity to the endoglin (ENG)-activin receptor-like kinase 1 (ALK1) receptor complex to control EC homeostasis and vascular quiescence *via* activation of Smad1/5/8 signaling^27–29^. The possible presence of a cross talk between OS and BMP9/10 signaling during AJ remodeling has not been addressed. In this work, the role of OS in VE-cadherin processing and AJ remodeling has been investigated and the involvement of MMPs and γ-secretase in this process has been assessed. Moreover, the role of the proteasome in VE-Cad/CTF2 metabolism has been evaluated. Indeed, proteasome activity plays a well-recognized role in the removal of CTFs derived from other γ-secretase substrates, such as several receptor tyrosine kinases^30^. Lastly, BMP9 and BMP10 have been tested for their effects on VE-cadherin processing and junction disassembly upon OS conditions.

## Results

### E-cadherin and VE-cadherin undergo similar processing in epithelial cells

Even though VE-cadherin is preferentially expressed in endothelial cells, it can be found in some cancer cells. We first tested the epithelial carcinoma cell line A431 that expresses both E-cadherin and VE-cadherin, in order to assess the similarities in the proteolytic cleavage of the two cadherins. We previously reported that E-cadherin processing by MMPs and γ-secretase can be induced in A431 cells upon treatments with the apoptosis inducer staurosporin (STS) and the calcium ionophore ionomycin (IONO)^11^. We found that cell exposure for 4-6 h to 1 μM STS, or for 30-90 min to 5 μM IONO, resulted in the cleavage of both E-cadherin (Fig.1a) and VE-cadherin (Fig.1b), and led to the generation of their respective ∼28 kDa C-terminal fragments, CTF2. These data show that apoptosis and calcium influx activate the proteolytic cleavage of both E- and VE-cadherins to generate a ∼28 kDa CTF2 fragment.

**Fig.1.**
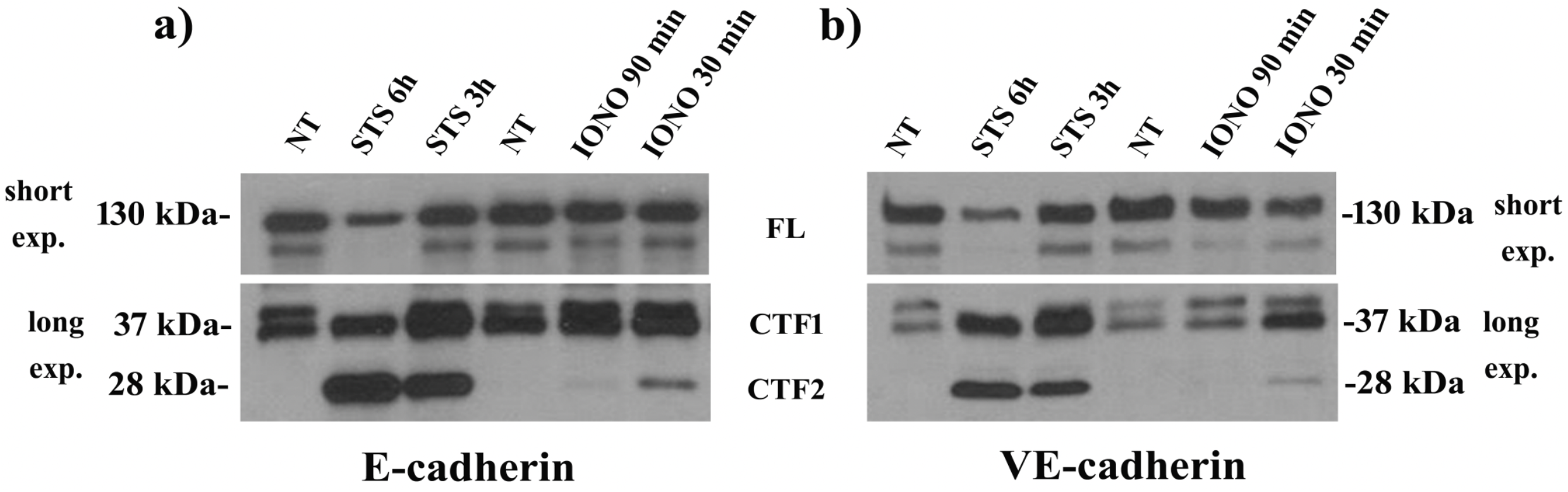
E- and VE-cadherin processing in epithelial cells. A431 cells were incubated for the indicated times in the absence (non-treated, NT) or presence of 1 μM staurosporin (STS) or 5 μM ionomycin (IONO). The blots show the expression of full-length (FL, 130 kDa upper bands), CTF1 (37 kDa middle bands) and CTF2 (28 kDa bottom bands) of E-cadherin (a) and VE-cadherin (b). Results are representative of 4 independent experiments. Different blot exposures were used in Figs. 1-3 and Supplementary Fig. 1 to facilitate the visualization of FL cadherins (short exposures, short exp.), as well as CTF1 fragments and CTF2 fragments (long exposures, long exp.), as indicated.

### Proteasome inhibition induces accumulation of the γ-secretase-derived VE-Cad/CTF2 fragment in endothelial cells

It has been demonstrated that γ-secretase products of other transmembrane proteins can be targeted by the proteasome^30^. The role played by the proteasome in degrading VE-Cad/CTF2 has been analyzed in HUVECs. As shown in Fig. 2a, in HUVECs treated with the proteasome inhibitor epoxomicin (EPOX, 1 μM, 3-6h), VE-Cad/CTF2 significantly accumulated, showing that VE-cadherin is produced at steady state in normal culture conditions and that the proteasome efficiently degraded VE-Cad/CTF2 in endothelial cells. HUVEC treatment with the γ-secretase inhibitor L-685,458 (GSI, 1 μM, 24h) fully blocked the accumulation of VE-Cad/CTF2 upon EPOX treatment, confirming that VE-Cad/CTF2 is produced by γ-secretase. GSI also led to the accumulation of the VE-Cad/CTF1 fragment, including in the absence of proteasome inhibition, indicating that (i) both VE-Cad/CTF1 and VE-Cad/CTF2 productions contribute to steady state VE-cadherin dynamics, (ii) VE-Cad/CTF2 is generated from the γ-secretase cleavage of VE-Cad/CTF1, and (iii) VE-Cad/CTF2 is efficiently cleared at steady state by the proteasome. Moreover, as shown in Fig. 2b, subcellular fractionation of EPOX-treated HUVECs showed that full-length VE-cadherin and VE-Cad/CTF1 mainly localized in membrane and cytoskeletal fractions, while VE-Cad/CTF2 was exclusively found in the cytosolic fraction, showing that VE-Cad/CTF2 is soluble and released into the cytosol upon γ-secretase cleavage.

**Fig.2.**
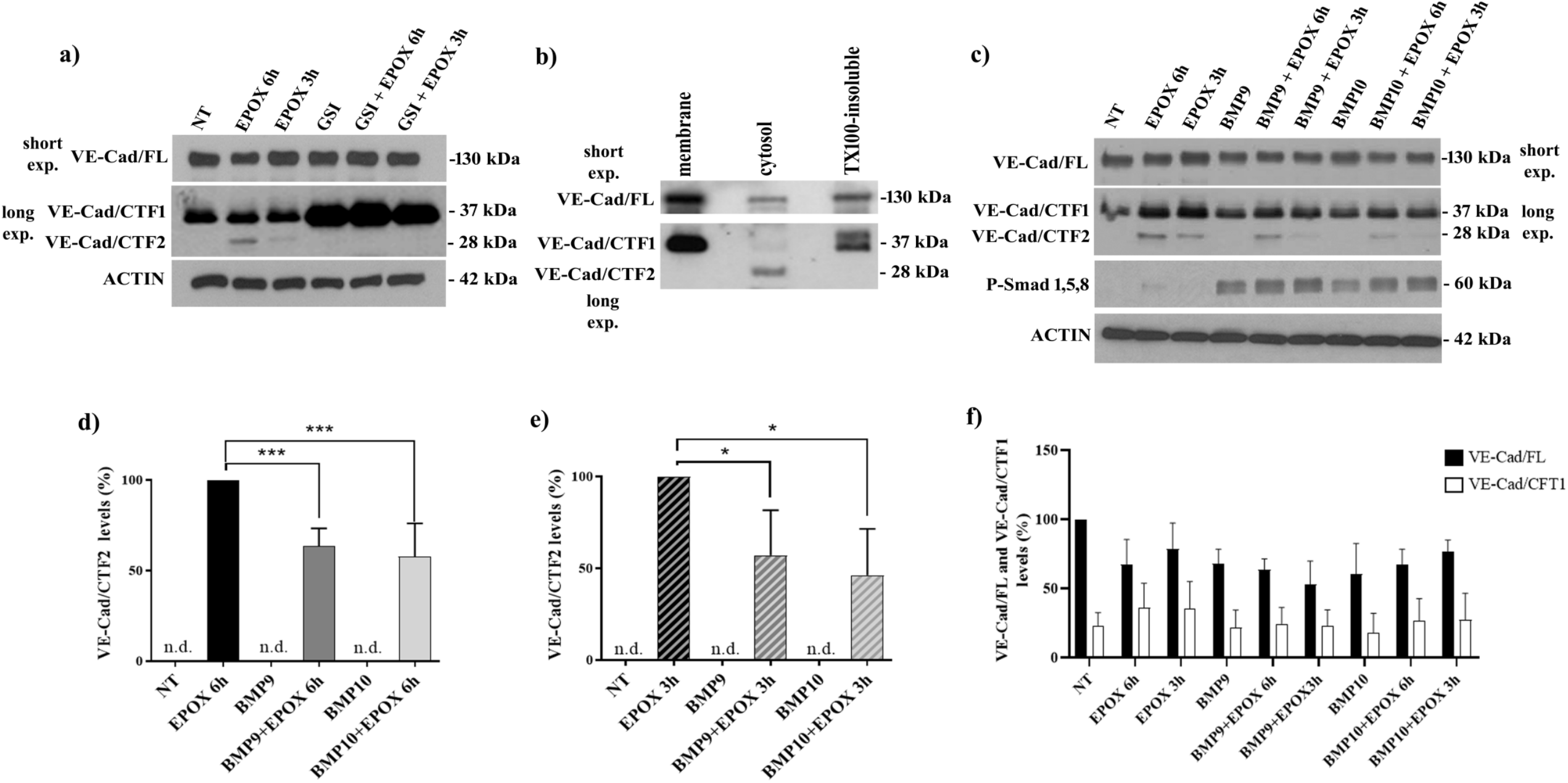
Role of γ-secretase, the proteasome, and BMP9/10 in the control of VE-Cad/CTF2 levels. a) HUVECs were pre-treated with GSI (L-685,458; 1 μM) for 24h and stimulated with epoxomicin (EPOX, 1 μM) for the indicated times. The blots show the expression level of VE-Cad/FL, VE-Cad/CTF1 and VE-Cad/CTF2 obtained from the same experiment (representative of 3 independent experiments). Actin was used as loading control. b) HUVECs treated for 6 h with EPOX were fractionated into membrane, soluble cytosolic, and Triton X-100-insoluble (TX100-insoluble) fractions, and the fractions obtained were probed by WB for VE-cadherin. c) HUVECs were exposed to BMP9 and BMP10 (10 ng/ml) for 24h before stimulation with EPOX (1 μM) for the indicated times. The blots show the expression level of VE-Cad/FL, VE-Cad/CTF1 and VE-Cad/CTF2 obtained from the same experiment (representative of 3 independent experiments). Membranes were reprobed for phospho-Smad1/5/8 (P-Smad1/5/8) to confirm BMP9/10 signaling activation. Actin was used as loading control. d and e) Densitometric analysis of VE-Cad/CTF2 levels in different experiments as in (c) (n=3). f) Densitometric analysis for VE-Cad/FL and VE-Cad/CTF1 levels in different experiments as above (n=3). Data in (d-f) are mean ± s.e.m, ***P < 0.005; *P < 0.05.

### BMP9/10 reduce VE-Cad/CTF2 levels in endothelial cells

To understand whether BMP9/10 signaling was involved in VE-cadherin processing, EPOX-treated HUVECs were stimulated with BMP9 or BMP10 (10 ng/ml, 24h). Both BMP9 and BMP10 significantly decreased VE-Cad/CTF2 levels in HUVECs exposed to EPOX for 3h or 6h (Fig.2c-2e). Full-length VE-cadherin and VE-Cad/CTF1 were not significantly affected by either EPOX or BMP9/10 treatments (Fig.2c and 2f). In these experiments, the levels of phospho-Smad1/5/8 - a transcriptional effector of BMP9/10 signaling - were analyzed and confirmed activation of the relevant signaling upon BMP9 and BMP10 treatments (Fig.2c). These findings suggest that BMP9/10 signaling negatively controls VE-Cad/CTF2 levels.

### Oxidative stress promotes VE-cadherin processing and VE-Cad/CTF2 production by ADAM10/17 and γ-secretase in endothelial cells

To study the role played by OS in VE-cadherin processing, HUVECs were exposed to increasing concentration of hydrogen peroxide (25-500 μM H_2_O_2_) for 6h. As shown in Supplementary Fig.1, WB analysis for VE-cadherin revealed the presence of a 28 kDa fragment in cells exposed to 500 μM H_2_O_2_ (Suppl. Fig.1 a). Moreover, the analysis of Annexin-V and PI positive cells performed on living cells showed that only the highest doses of H_2_O_2_ used was able to significantly increase the number of apoptotic cells (Suppl. Figs.1b and c), suggesting that the formation of the VE-Cadherin 28kDa fragment is linked to OS-dependent cell damage. To analyze the involvement of MMPs and γ-secretase in the formation of the identified H_2_O_2_-induced VE-cadherin 28kDa CTF, HUVECs were pretreated with 1 μM GSI or 10 μM GI254023X (ADAM17 and ADAM10 inhibitor) and then exposed to 500 μM H_2_O_2_. As shown in Fig.3a, H_2_O_2_-induced 28kDa CTF production was fully blocked by GSI, confirming that this fragment is VE-Cad/CTF2. VE-Cad/CTF2 production induced by H_2_O_2_ was also prevented by the MMP inhibitor. Of note, while GSI prevented VE-Cad/CTF2 formation and favored the accumulation of VE-Cad/CTF1, the inhibitor of MMPs was able to prevent VE-Cad/CTF1 formation highlighting that the two enzymes act sequentially. Indeed, MMPs was involved in the cleavage of full-length VE-cadherin generating VE-Cad/CTF1, and γ-secretase cleaved VE-Cad/CTF1, generating VE-Cad/CTF2.

**Fig.3.**
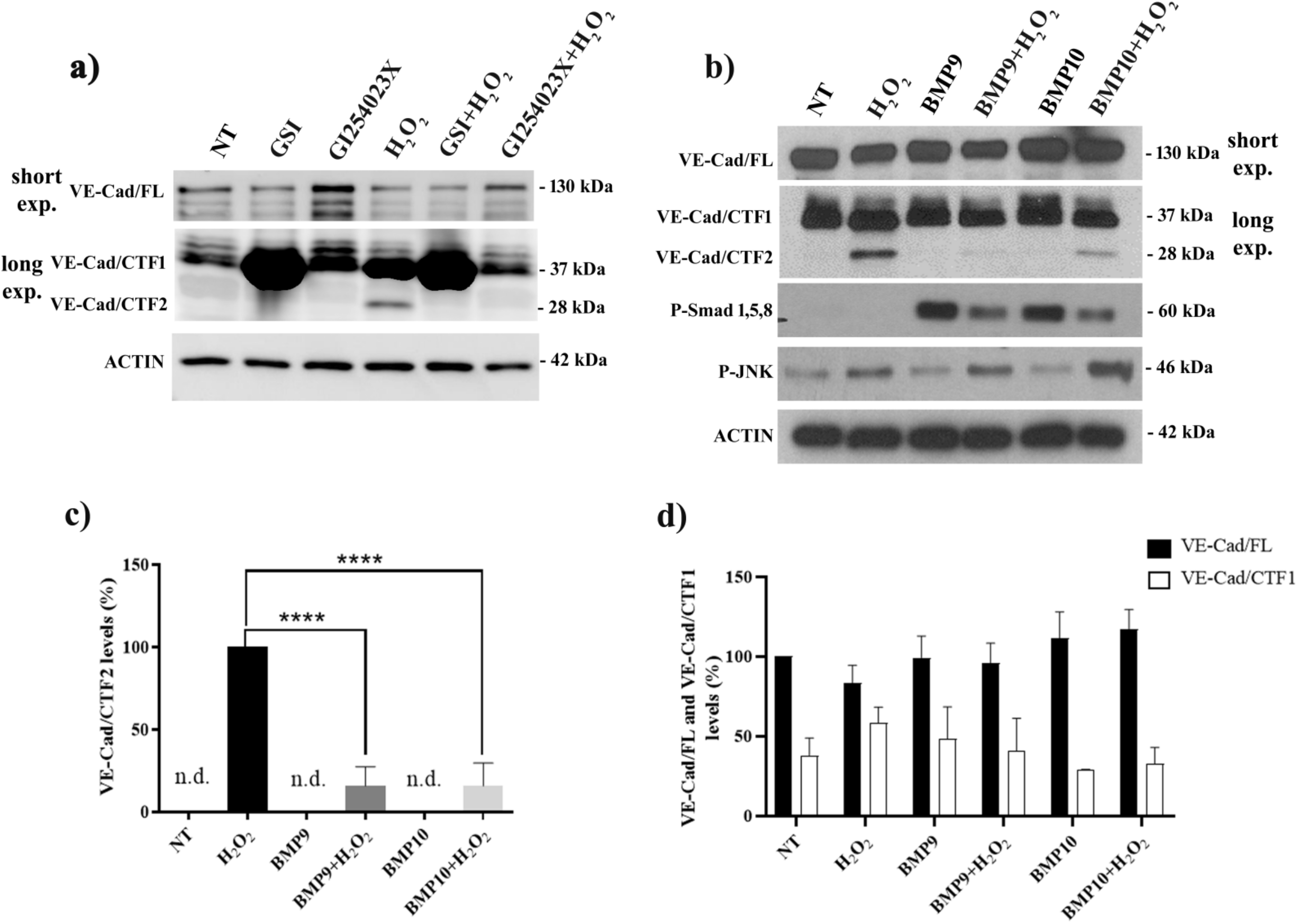
Role of γ-secretase, the metalloproteases, and BMP9/10 in VE-cadherin processing under oxidative stress. a) WB analysis of VE-cadherin in HUVECs exposed to GSI (L-685,458; 1 μM) or GI254023X (10 μM) for 24h before stimulation with H_2_O_2_ (500 μM) for 6h. Actin was used as loading control. Note that a digital imager was used for this analysis and some CTF1 bands were overexposed to allow detection of CTF2, see also the corresponding uncropped scans in supplementary material. b) WB analysis of VE-cadherin in HUVECs exposed to BMP9 and BMP10 (10 ng/ml) for 24h before stimulation with H_2_O_2_ (500 μM) for 6h. Phospho-Smad1/5/8 was used as a BMP9/10 signaling activation marker. Results are representative of 3 independent experiments. c) Densitometric analyses of VE-Cad/CTF2 levels in different experiments as in (b). d) Densitometric analysis of VE-Cad/FL and VE-Cad/CTF1 levels in different experiments as in (b). Data in (c and d) are mean ± s.e.m. of 3 independent experiments, ****P<0.005.

### BMP9/10 reduce oxidative stress-induced VE-Cad/CTF2 levels

Considering our results showing an effect of BMP9/10 on VE-Cad/CTF2 accumulation in HUVEC treated with the proteasome inhibitor, we asked whether these ligands also control VE-Cad/CTF2 levels upon OS. As shown in Fig.3b, pretreatment with 10 ng/ml BMP9 or BMP10 for 24h significantly reduced VE-Cad/CTF2 levels induced by 500 μM H_2_O_2_ for 6h. Densitometric analysis showed ∼80 % reduction of VE-Cad/CTF2 levels by BMP9 or BMP10 in cells treated with H_2_O_2_ (Fig.3c), while full-length VE-cadherin and VE-Cad/CTF1 levels were unaffected by H_2_O_2_ or BMP9/10 treatments (Fig.3d). As expected, phospho-Smad1/5/8 levels increased upon BMP9 and BMP10 treatments, and phospho-JNK, a marker of OS, accumulated upon H_2_O_2_ stimulation (Fig.3b). Thus, BMP9/10 signaling potently repressed VE-Cad/CTF2 accumulation upon OS. Of note, BMP9 treatment failed to prevent the H_2_O_2_-mediated increase of endothelial permeability assessed in a transwell assay (Supplementary Fig.2), suggesting that γ-secretase processing of VE-cadherin is not involved in this process.

### Oxidative stress-induced redistribution of the VE-cadherin/β-catenin/F-actin complex is partially prevented by BMP9/10 treatment or MMPs inhibition

Since cadherin processing modifies its binding to β-catenin^11^, we then analysed by immunofluorescence the co-localization of VE-cadherin and β-catenin in HUVECs treated with H_2_O_2_. In particular, to understand the effect of OS on the distribution of other AJ components, a triple staining for VE-cadherin, β-catenin and F-actin was performed. As shown in Supplementary Fig.3, a VE-cadherin typical zip-line membrane localization was observed in untreated cells; β- catenin closely followed VE-cadherin distribution and F-actin was visible inside cells. HUVECs treated with 25 μM-100 μM H_2_O_2_ did not show significant changes in VE-cadherin, β-catenin and F-actin subcellular localization, with a colocalization of VE-cadherin and β-catenin that was maintained at the cell surface. However, clear cell-cell contact modifications were visible, suggesting that permeability was affected at concentration as low as 25 μM. Actin fibers were well visible in the cytosol and some discrete colocalization with VE-cadherin and β-catenin could be observed (white punctas in merged panels). Cells treated with 200 μM H_2_O_2_ showed cell-cell contact defects, but some colocalization between VE-cadherin and β-catenin was still present, and F-actin was well defined inside the cells. 500 μM H_2_O_2_ induced the strongest effect by reducing VE-cadherin localization at the membrane. Some VE-cadherin colocalization with β-catenin was still present, but F-actin was strongly disorganized and its colocalization with β-catenin and VE-cadherin was observed in discrete areas. The effect of BMP9/10 pre-treatments and MMPs inhibition were analysed in cells exposed to 500 μM H_2_O_2_. As shown in Fig.4, cell pretreatment with 10 ng/ml BMP9 or BMP10 prevented the reduction of VE-cadherin expression at the cell surface, reduced VE-cadherin/β-catenin redistribution, and limited F-actin disorganization due to H_2_O_2_. Moreover, cell exposure to GI254023X prevented VE-cadherin/β-catenin redistribution induced by 500 μM H_2_O_2_ (Fig.4). However, the efficacy in preventing F-actin disorganization induced by H_2_O_2_ was limited, indicating that other additional mechanisms implicating MMP- and γ- secretase-independent pathways contributed to VE-cadherin redistribution. Of note, HUVECs exposure to 10 ng/ml BMP9 or BMP10 alone showed slight modifications of VE-cadherin, β- catenin and F-actin staining that accompanied possible changes in AJ integrity, while GI254023X *per se* did not induce any changes (Supplementary Fig.4).

**Fig.4.**
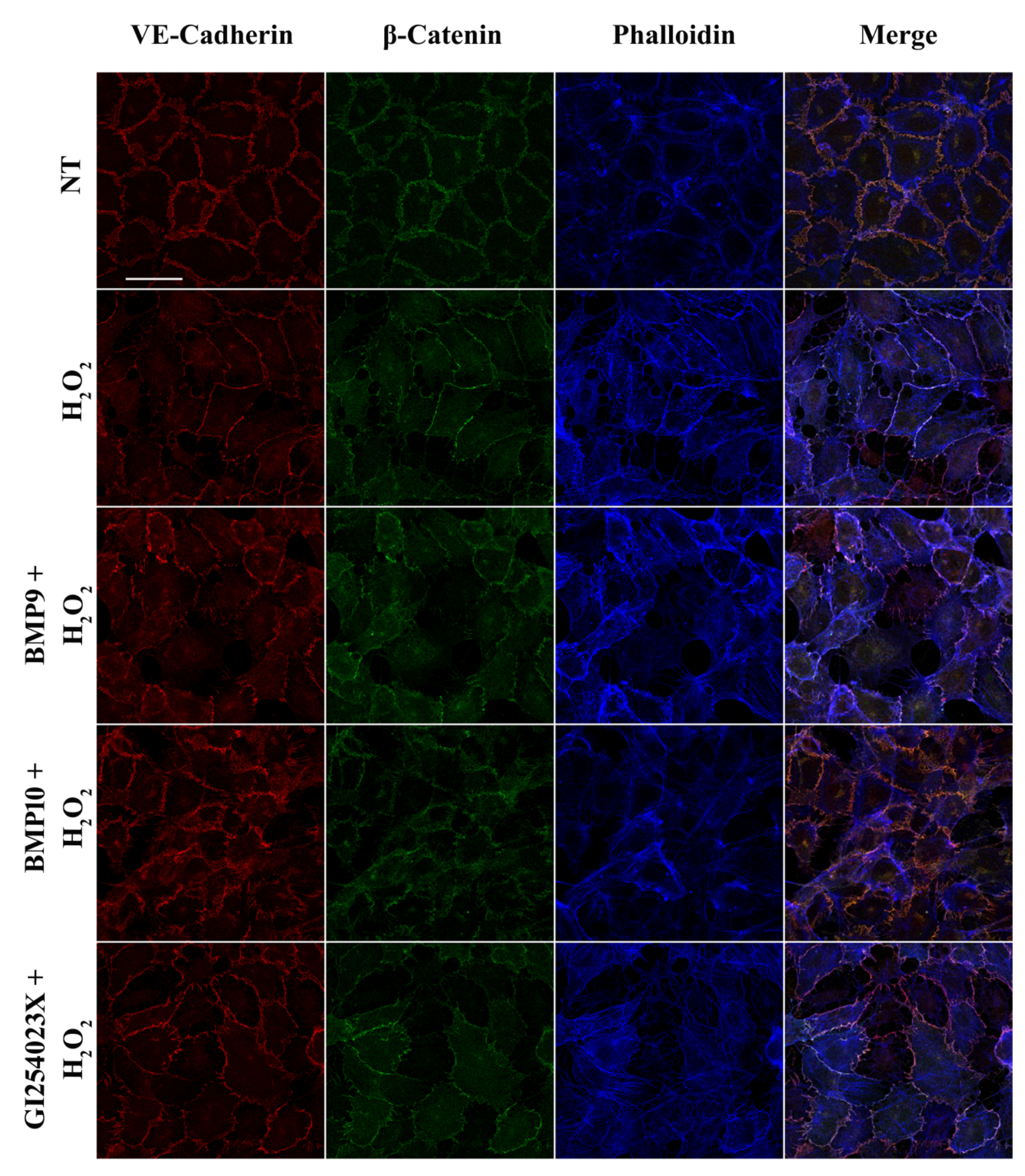
VE-cadherin, β-catenin and F-actin distribution in HUVECs exposed to oxidative stress. HUVECs were pre-treated with BMP9, BMP10 (10 ng/ml) or GI254023X (10 μM) for 24h and then exposed to H_2_O_2_ (500 μM) for 6h. HUVECs were stained for VE-Cadherin (red), β-Catenin (green) using specific antibodies. Phalloidin was used to stain F-actin (blue). Scale bar=30 μm.

### F-actin redistribution upon oxidative stress does not implicate actin cytoskeleton depolymerization

To investigate whether F-actin modification due to OS could be related to a depolymerization of the actin cytoskeleton, the effect of H_2_O_2_ was compared to the effect of a classic depolymerizing agent, cytochalasin B. As shown in Fig.5a, F-actin/G-actin ratio (measured by WB analysis) decreased as expected in cells exposed to cytochalasin B, but not in cells exposed to H_2_O_2_, where the ratio actually increased significantly. Immunofluorescence analysis of F-actin confirmed the different effects obtained with the two treatments. Indeed, F-actin appeared redistributed inside the cells after H_2_O_2_ treatment, while in cells exposed to cytochalasin B, F-actin appeared fragmented, hence disassembled (Fig.5c). Moreover, F-actin fluorescence quantification showed no significant changes in cells treated with H_2_O_2_ compared to untreated cells, while a strong reduction was observed after cytochalasin B treatment (Fig.5b).

**Fig.5.**
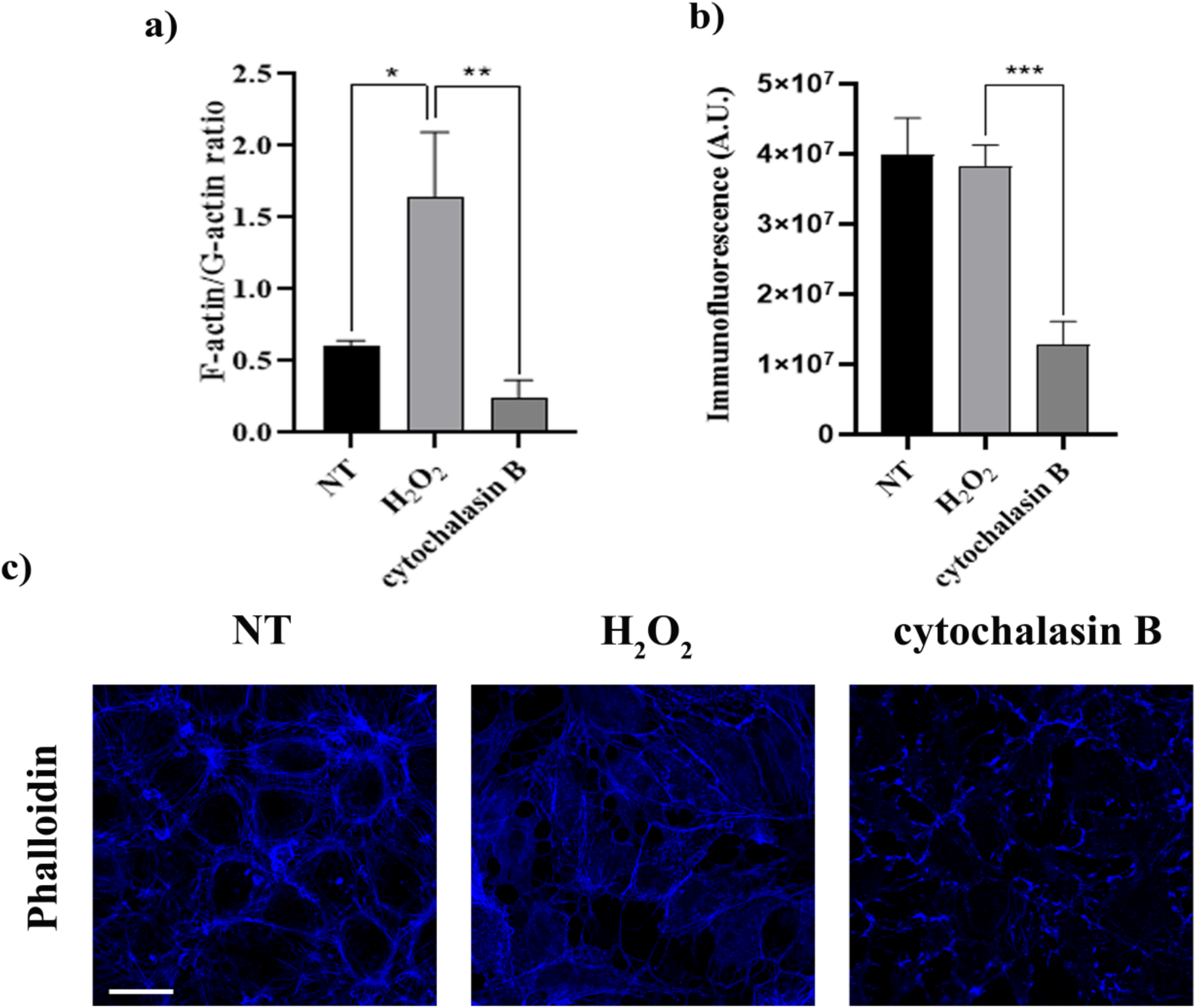
F-actin modification due to oxidative stress. a) F-actin/G-actin ratio was measured by WB analysis in HUVECs stimulated with H_2_O_2_ (500 μM) for 6h or cytochalasin B (5 μM) for 10 min. Data represent the mean ± s.e.m. of 6 independent experiments, *P<0.05; **P<0.01; b and c) After HUVEC treatment with H_2_O_2_ (500 μM) for 6h and cytochalasin B (5 μM) for 10 min, a Phalloidin staining (F-actin marker) was performed. Panels in (c) show one representative experiment. Scale bar=30μm. F-actin immunofluorescence was quantified by ImageJ-win64. Each measurement included twelve areas of interest per image. A.U., arbitrary units. Data in (b) are mean ± s.e.m. of 3 independent experiments, ***P<0.001.

### BMP9/10 pretreatments reduce oxidative stress-induced apoptosis, favoring endothelial cells recovering

To investigate the contribution of BMP9/10 in endothelial homeostasis, we analyzed the effect of BMP9 and BMP10 pretreatments on the apoptosis induced by a 6h exposure to 500 μM H_2_O_2_ followed by 24h of recovery. As shown in Fig. 6, the percentage of Annexin V positive cells were significantly reduced by BMP9 and BMP10 pretreatments following H_2_O_2_ exposure.

**Fig.6.**
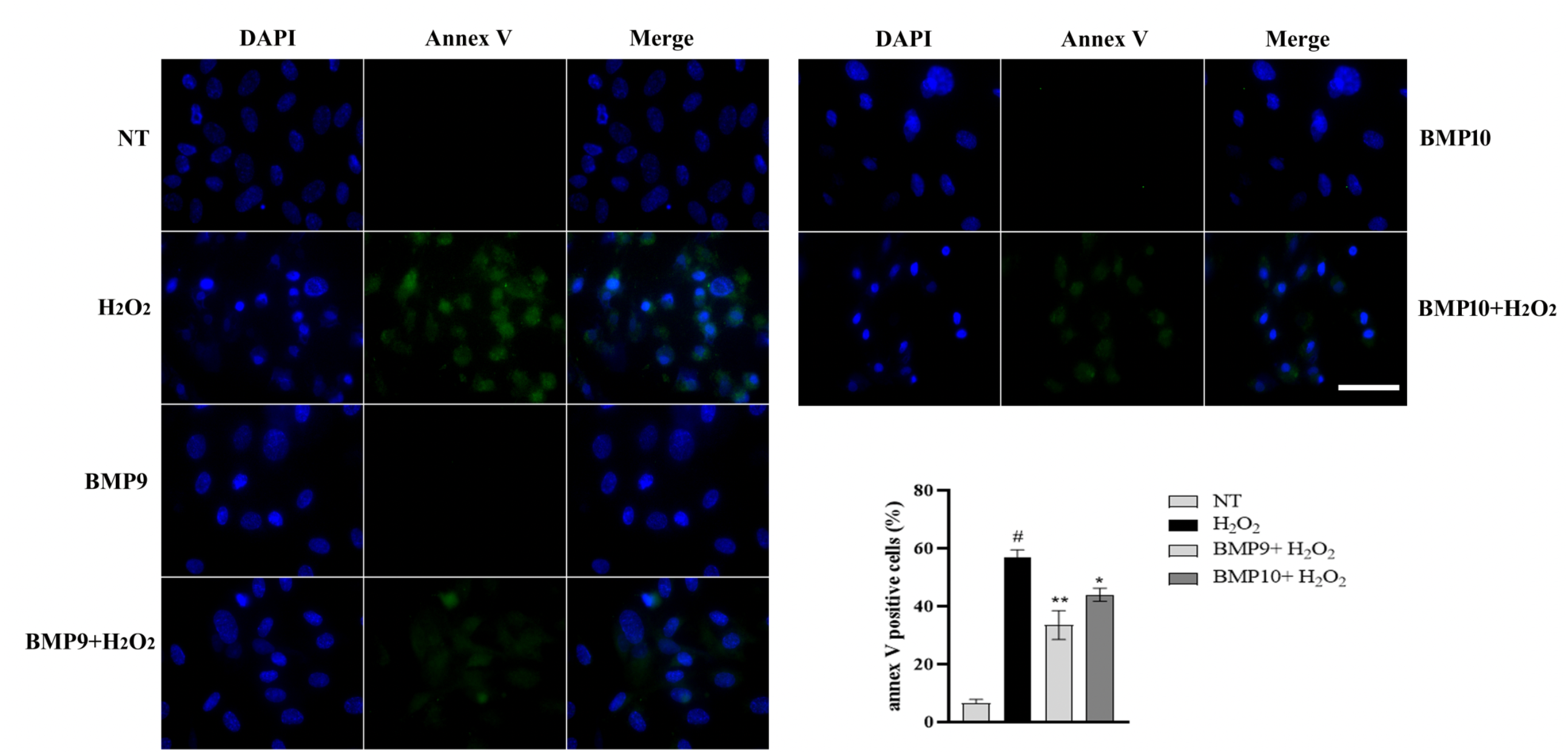
BMP9 and BMP10 mitigate apoptosis induced by H_2_O_2_. Annexin (Annex) V-FITC staining analysis (green) in HUVECs following a 6h treatment with 500 μM H_2_O_2_ in the absence or presence of a 24 h pre-incubation with BMP9 and BMP10 (10 ng/mL). Nuclei are counterstained by using DAPI, as detailed in Materials and Methods. Scale bar=50 μm. Histogram shows densitometric analysis of Annexin V-positive cells (%) treated as above (n=3). Data are mean ± s.e.m, # P<0.001 vs. NT; **P<0.01, *P<0.05 vs. H_2_O_2_.

## Discussion

In this work, we show for the first time that VE-cadherin undergoes an MMP- and γ-secretase-dependent processing, which is enhanced by OS and is modulated by both the proteasome and BMP9/10 signaling. In endothelial cells, AJs play a crucial role in maintaining barrier integrity^31^ and regulating permeability^32^ and angiogenesis^33^, and VE-cadherins, the core components of the AJs, control survival signals^34^. Thus, AJ disassembly and VE-cadherin modifications have been well demonstrated during endothelial cell damage^34,35^ in different pathophysiological conditions^36^ but the proteolytic pathways involved in VE-cadherin processing remain only partially understood.

By using an epithelial cell line (A431) expressing both E-cadherin and VE-cadherin, we demonstrated that apoptotic induction or intracellular Ca^2+^ imbalance favors the cleavage of the two types of cadherins, with the generation of two C-terminal fragments, namely CTF1 and CTF2. The proteolytic processing of E-cadherin was already known, and our results are in agreement with previous studies that reported E-cadherin endoproteolysis upon apoptosis and calcium influx to generate the E-Cad/CTFs^11,37^. However, we provide here the first evidence of VE-Cad/CTF2 detection, although it existence was already indirectly demonstrated upon inhibition of γ-secretase and accumulation of its substrate VE-Cad/CTF1^12^.

To better understand the mechanisms of VE-cadherin processing regulation, we analyzed the involvement of the proteasome in VE-Cad/CTF2 clearance. The role played by the proteasome in the clearance of γ-secretase-produced CTFs has been demonstrated for IL-11R, CSF1R, IGF1R, MET, or TIE1^30,38^. Interestingly, we found that the inhibition of proteasomal activity using epoxomicin has a strong stabilizing effect on VE-Cad/CTF2 levels in HUVECs. To the best of our knowledge, this is the first evidence proving the involvement of the proteasome in the degradation of VE-Cad/CTF2 in primary endothelial cells. Notably, we also provided evidence that VE-Cad/CTF2 was soluble and released in the cytosol, consistent with what was highlighted previously for E-Cad/CTF2^11^.

Next, we investigated the role of OS in VE-cadherin processing. Endothelial pathophysiology can be driven by OS^39^. Indeed, OS is involved in modulating endothelial functions, such as vascular permeability^40^, and its contribution to several endothelial-related pathologies from diabetes to atherosclerosis has been demonstrated^18,19^. However, the role played by OS in AJ disassembly is still poorly understood. Our data demonstrate that H_2_O_2_ treatment was able to favor VE-Cad/CTF2 formation and to modify VE-cadherin, β-catenin and F-actin interactions. Indeed, under OS, the typical VE-cadherin staining on cell membrane was lost with a change in junction conformation from zip to straight. The straight conformation has been already observed when the cells are exposed to different flow patterns or VE-cadherin is blocked with specific antibody^31,41^.

We further observed that F-actin is strongly modified and undergoes an evident disorganization with an increase in F-actin/G-actin ratio upon OS. OS has already been shown to induce changes in actin cytoskeleton structure in endothelial cells^42,43^ and myoblast^44^, and to lead to an increased in F-actin/G-actin ratio, which was proposed to be due to oxidative inhibition of cofilin^45^. Further studies will be required to determine whether a similar mechanism can apply to our experimental condition and might be regulated by BMP9/10-dependent VE-cadherin proteolysis. Importantly, MMP inhibition obtained using GI254023X, which is highly selective for ADAM10 and ADAM17^46^, significantly prevented VE-Cad/CTF1 and VE-Cad/CTF2 formation, as well as VE-cadherin redistribution induced by H_2_O_2_. Our data are in agreement with other papers in the literature demonstrating the upstream involvement of MMPs in VE-cadherin shedding and proteolytic cleavage during apoptosis^7,12^. Besides, other studies have demonstrated the strong correlation between OS and MMP activation in the context of age-related macular degeneration (AMD)^47^ and in coronary artery disease^48^.

Interestingly, the analysis of the interconnection between VE-cadherin, β-catenin and actin showed the ability of the MMP inhibitor to prevent VE-cadherin/β-catenin redistribution induced by H_2_O_2_, while the effect on actin redistribution was limited. This result suggests that AJ reorganization upon OS is both MMP/γ-secretase-dependent (VE-cadherin processing and VE-cadherin/β-catenin redistribution) and MMP-independent (F-actin belt disorganization). Additional investigation will be required to further our understanding of the broader consequences of OS-triggered actin cytoskeleton deregulation for AJ disassembly.

Previously, we demonstrated that E-cadherin proteolytic degradation was dependent on MMPs and γ-secretase^11^. The role of γ-secretase was also highlighted by other studies showing that the γ-secretase complex is incorporated into the E-cadherin-catenin complex under conditions promoting cell adhesion^49,50^ while under conditions of cell-cell dissociation or apoptosis the activity of γ-secretase promotes AJ dissociation^11^. The activation of γ-secretase in response to OS was shown for the proteolytic processing of APP, a protein involved in Alzheimer’s disease^51,52^. In this study, we have shown that a γ-secretase inhibitor prevented VE-Cad/CTF2 accumulation in the presence of epoxomicin (that blocks VE-Cad/CTF2 degradation) or H_2_O_2_ (that increases VE-Cad/CTF2 levels) to lead to an accumulation of VE-Cad/CTF1. Our data points therefore to a two steps proteolytic cleavage of full-length VE-cadherin in response to OS that sequentially involves metalloproteases (ADAM10 and ADAM17) and γ-secretase.

Loss of function mutations in ALK1 and ENG cause hereditary hemorrhagic telangiectasia (HHT), a genetic bleeding disorder characterized by EC proliferation and migration defects that leads to the development of systemic vascular lesions^53–56^. VE-cadherin’s role in AJ disassembly and angiogenesis could be relevant to HHT pathology^57^. Notably, ROS levels were reported to increase because of uncoupled eNOS formation in different tissues of ENG and ALK1 heterozygous knockout mice^58,59^. In this context, our data show that BMP9/10 signaling could reduce steady-state VE-Cad/CTF2 levels upon OS. In future work, it will be interesting to determine in HHT models whether VE-cadherin proteolytic processing accompanies vascular lesion development and is a consequence of reduced BMP9/10-ALK1-ENG signaling, as suggested by our in vitro studies. Moreover, BMP9 and BMP10 were able to reduce VE-cadherin/β-catenin redistribution and actin modification in the presence of H_2_O_2_. Whether this effect is due to a reduction of VE-cadherin cleavage has to be elucidated. The existence of a cross talk between VE-cadherin processing and TGF-β receptor signaling^58^ or BMP signaling has been reported before. For instance, it has been shown that VE-cadherin can associate with ALK2 and BMPRII in a BMP6-dependent manner and that BMP6 induces EC permeability changes via the promotion of VE-cadherin internalization and phosphorylation at Tyr-685^60^. Knowing that ALK2 and BMPRII can also bind to BMP9, and BMPRII is a type II receptor candidate for the ALK1 complex in endothelial cells^61^, a possible mechanism of VE-cadherin processing regulation by BMP9 might implicate ALK2 and/or BMPRII binding regulation to VE-cadherin and the resulting trafficking changes of the cell-cell adhesion protein. Additional work will be required to clearly address these possible mechanisms.

Finally, we demonstrated that BMP9 and BMP10 exert pro-survival effects in HUVECs by reducing OS-dependent apoptosis and facilitating cell recovering. This result is in agreement with other report on the protective effect of BMP9 on TNF-α-induced apoptosis of pulmonary ECs^62^. We also showed that BMP9 is not able to reduce OS-dependent hyperpermeability, excluding the possibility that the BMPs promote survival upon OS *via* this mechanism. Additional investigation will be needed to clarify the exact mechanism involved in the pro-survival effect of BMP9/10 during oxidative conditions. BMP9/10 signaling controls several transcriptional responses and some of these pathways might contribute to the observed protective effect against apoptosis and VE-cadherin processing upon H_2_O_2_ challenge. It is also important to note that VE-Cad/CTF2 itself could trigger specific signal transduction pathways during oxidative stress, and via some of these pathways, could interact with BMP9/10 signaling. Therefore, it will be interesting to determine whether these signaling cross talks are involved in the presented mechanism.

In sum, our work sheds light on a mechanism of AJ reorganization upon OS, which implicates BMP9/10 signaling in VE-cadherin processing regulation, a previously unsuspected function of BMP9/10 with possible relevance for vascular diseases such as HHT or pulmonary arterial hypertension.

## MATERIALS AND METHODS

### Cell Lines and treatments

Epithelial carcinoma cell line A431 (American Type Culture Collection) was grown in Dulbecco’s Modified Eagle’s medium (DMEM, Thermo Fisher Scientific, USA) supplemented with 10 % Fetal Bovine Serum (FBS, Thermo Fisher Scientific), 5 % penicillin/streptomycin solution (Thermo Fisher Scientific), 2 mM glutamine (Sigma-Aldrich, USA). Cells were subcultured every 2 days, maintained in 5 % CO_2_ at 37° C and used between passage 2 and 15. HUVECs (National Cord Blood Program at Northwell Health^63^ and Core Facility of IRCCS AOU San Martino, Genoa, Italy) were routinely cultured in Endothelial Cell Medium (ECM, Clinisciences, Italy) supplemented with 5 % Fetal Bovine Serum (FBS, Clinisciences), 1 % Endothelial Cell Growth Supplement (ECGS, Clinisciences) and 1 % penicillin/streptomycin solution (Clinisciences). Cells were subcultured every 4 days, maintained in 5 % CO_2_ at 37°C and used between passage 2 and 11. A431 cells were exposed to 1 μM staurosporine (Cayman chemical, USA) or 5 μM ionomycin (Cayman chemical). HUVECs were exposed to 500 μM H_2_O_2_ (Sigma-Aldrich) or 1 μM epoxomicin (Cayman chemical). Samples were treated with 10 μM GI254023X (Cayman chemical), 1 μM L-685,458 (GSI, Sigma-Aldrich, USA), 10 ng/ml recombinant human BMP9 and BMP10 (R&D Systems), or 5 μm cytochalasin B (Sigma-Aldrich).

### Endothelial Permeability Assay

HUVECs permeability was assessed by using Endothelial Transwell Permeability Assay Kit (Cell Biologics, Chicago, USA) following manufacturer instructions. Briefly, cells were seeded on the 6.5 mm transwell insert membrane in 200 μl ECM medium, and transferred to a 24-well plate containing 1 ml of ECM media. Cells were treated at confluence and 5 μl of streptavidin-HRP were added to each well. At the end of the treatments, absorption of the lower chamber media was measured at 450 nm in 96-well ELISA plates.

### Viability assay

Viability has been measured by Trypan Blue exclusion test (Sigma-Aldrich). Cell viability was calculated using the ratio of viable/total cells and expressed as percentage values.

### Apoptosis detection by immunofluorescence assay

To assess apoptosis in living cells, HUVECs were seeded in 8-well chamber slide (IBIDI®, Gräfelfing, Germany) at the density of 100×10^3^ cells/well. Apoptosis detection Kit (Dojindo Molecular technology Inc. Rockville, MD, USA) was used according to the manufacturer’s protocol. Briefly, at the end of treatments cells were washed, and incubated with FITC-Annexin V (1:20) plus propidiun iodide (PI, 1:20) in the supplied buffer for 10 minutes. Then, cells were washed and immediately observed using a three-channel TCS SP2 laser-scanning confocal microscope (Leica, Wetzlar, Germany), equipped with 458, 476, 488, 514, 543 and 633 nm excitation lines.

To acquire images also of fixed cells, FITC-Annexin V Apoptosis Detection Kit (BioVision Research, USA) was used according to the manufacturer’s protocol. HUVECs were seeded on 8-well chamber slides (Nalge Nunc International) at the density of 100×10^3^ cells per well. At the end of the treatments, cells were incubated for 10 min in the dark with Annexin V-FITC (1:100) and then with DAPI solution (1:1000) for the same time. Subsequently, HUVECs were fixed in 2 % PFA before visualization. Images were acquired using epifluorescence microscope (Olympus 1×81) with a 60x oil immersion objective.

### F-actin/G-actin extraction

At the end of the treatments, cells were washed with cold PBS and harvested in lysis buffer (10 mM HEPES, pH 7.0, 50 mM NaCl, 1% Triton X-100, 1 mM MgCl_2_, 2.5 mM EGTA). After a 10 min incubation on ice, the samples were centrifuged at 16,000 x g for 10 min and the supernatants (G-actin fraction) were collected. The pellets were washed once with lysis buffer and resuspended in the same volume as the supernatants to obtain the F-actin fraction.

### Preparation of total cell lysates

Total protein extraction was performed using Laemli Buffer 1X (1 M Tris-HCl pH 6.8, 35 % Glycerol, 20 % SDS, 1 % Blue bromophenol, 7 % β-Mercaptoethanol). Protein content was measured using BCA test (Pierce BCA Protein Assay Kit, Thermo Scientific, Rockford, USA).

### Subcellular fractionation

At the end of the treatments, confluent HUVEC cells (in one 100-mm dish) were rinsed and scraped in hypertonic buffer A (0.25 M sucrose, 5 mM EDTA, 10 mM β-mercaptoethanol, 10 mM Hepes pH 7.5, supplemented with 1× Complete protease inhibitor cocktail, Roche, Germany), passed through 27 G syringe and left 20 min in ice. The lysate was centrifuged at 100,000 g for 30 min at 4°C to separate the cytosolic and crude membrane fractions. The pellet (crude membrane fraction) was suspended in buffer A containing 0.3 % Triton X-100. The suspension was incubated 20 min in ice and then centrifuged at 100,000 g for 30 min at 4°C to separate the membrane and cytoskeleton (Triton X-100-insoluble fractions). The pellet (Triton X-100-insoluble fraction) was solubilized by sonication in RIPA buffer, left 20 min in ice and then centrifuged at 14,000 g for 15 min at 4°C.

### Immunoblotting

Total protein lysates were subjected to electrophoresis on home-made acrylamide gels and Western blotting, following standard protocol. Immunodetection was performed using VE-cadherin mouse monoclonal antibody (mAb) sc-9989 IgG_1_ (F-8, C-terminal epitope; Santa Cruz Biotechnology, USA), E-cadherin mouse mAb (BD Biosciences, USA), phospho-Smad1 (Ser463/465) / Smad5 (Ser463/465) / Smad9 (Ser465/467) (D5B10, Cell Signaling Technology, USA) rabbit mAb and phospho-SAPK/JNK (Thr183/Tyr185) (G9, Cell signaling) mouse mAb, and specific secondary antibodies (anti rabbit/anti mouse IgG-HRP, Euroclone S.p.A, Italy). The blotting membranes were reprobed with loading control antibodies, mouse anti-actin (Santa Cruz), or mouse anti-tubulin (Abcam, UK). Protein expression was determined by enhanced chemiluminescence (ECL, Advansta, USA) and analyzed by densitometry using ImageJ studio Lite software.

### Confocal immunofluorescence

For confocal analysis HUVECs were seeded on 8-well chamber slides (Nalge Nunc International, USA) at the density of 100 × 10^3^ cells per well. At the end of the treatments, cells were fixed in 4 % PFA (paraformaldehyde) for 15 min, washed in PBS, permeabilized with 1 % Triton X-100 + 1 % BSA for 5 min, and incubated with primary antibody overnight at 4°C. Anti-VE-cadherin (mouse, Santa Cruz Biotechnology, 1:100) and anti-β-catenin (rabbit, Cell Signaling Technology, 1:100) antibodies were used followed by specific secondary Alexa Fluor antibodies (1:500 anti-mouse Alexa Fluor 546 and 1:500 anti-rabbit Alexa 488 Thermo Fisher). For actin staining, Phalloidin-iFluor 633 reagent was used (Abcam, 1:1000). Nuclei were stained with To-Pro 3 iodide (1 μg/ml), or propidium iodide (1 μg/ml). Slides were mounted, and images were acquired using a 3-channel TCS SP2 laser-scanning confocal microscope (Leica Mycrosystems, Germany) and STELLARIS 8, Falcon FLIM confocal (Leica Microsystems).

### Statistical analysis

All data denote the mean ± S.E.M of different experiments, as indicated. Statistical analyses were performed using Prism software package (GraphPad software, San Diego, CA). One-way analysis of variance (ANOVA) and Dunnett’s multiple comparison tests were applied when comparing more than three groups, with p< 0.05 considered significant.

## Abbreviations

A431: Epidermoid carcinoma human cell line
AVMs: Arteriovenous Malformations
AJ: Adherens junctions
BMP: Bone Morphogenetic Protein
CTF1: C-terminal fragment 1
CTF2: C-terminal fragment 2
EPOX: Epoxomicin
FL: Full Length
H_2_O_2_: Hydrogen Peroxide
HHT: Hereditary Hemorrhagic Telangiectasia
HUVECs: Human Umbilical Vein Endothelial Cells
MMPs: Matrix Metalloproteinases
OS: Oxidative stress
TGF: Transforming Growth Factor
VE-cadherin: Vascular Endothelial Cadherin

## Acknowledgements

We thank Pallavi Chandakkar and Haitian Zhao (The Feinstein Institutes for Medical Research) for excellent technical assistance. We are also grateful to Elena Gatta and the Nanoscale Biophysics Group at DIFILAB, Department of Physics, University of Genoa for their support and assistance in the use of Stellaris 8, Tau-STED, Falcon FLIM, Leica Microsystems Confocal microscope.

## Author contributions statement

C.I., M.N. and P.M. conceptualized the study; C.I., M.P., S.R., C.N.M. and P.K.C. developed methodology; C.I., M.P., A.L.F. and C.D. performed the experiments; C.I., M.N. and P.M. wrote the manuscript. All authors reviewed the manuscript.

## Additional information

The authors declare no competing interests.

## Figure legends

**Supplementary Fig.1: VE-Cad/CTF2 generation and apoptosis induction in HUVECs exposed to increasing concentration of H**_**2**_**O**_**2**_. a) WB analysis of VE-cadherin in HUVECs treated with increasing concentration of H_2_O_2_, as indicated. Actin was used as loading control. Results are representative of 2 independent experiments. b) Percentage of Annexin V FITC positive cells and propidium iodide (PI) positive cells was calculated counting 4 fields for each experimental condition in 3 independent experiments. Data are mean ± s.e.m, *P<0.001 vs. AnnexV FITC positive cells in NT, 100μM and 200 μM H_2_O_2_, # P<0.001 vs. PI positive cells in NT, 100 μM and 200 μM H_2_O_2_. c) Representative images of HUVECs exposed to increasing concentrations of H_2_O_2_ and stained with annexin V-FITC/PI, as described in materials and methods. Scale bar=50 μm.

**Supplementary Fig 2: BMP9 does not prevent oxidative stress-induced permeability**. The graph shows permeability of HUVECs pre-treated with BMP9 and exposed to 500 μM H_2_O_2_ for 6 hours. Data are mean ± s.e.m. of 3 independent experiments. *P<0.01 vs NT.

**Supplementary Fig.3: VE-cadherin, β-catenin, and F-actin distribution in HUVECs exposed to increasing concentration of H**_**2**_**O**_**2**_. HUVECs were exposed to H_2_O_2_ (25-500 μM) for 6h. HUVECs were stained for VE-Cadherin (red), β-Catenin (green) using specific antibodies. Phalloidin was used to stain F-actin (blue). Scale bar = 30 μm

**Supplementary Fig.4: VE-cadherin, β-catenin, and F-actin distribution in HUVECs exposed to BMP9/10 or MMPs inhibitor**. HUVECs were pre-treated with BMP9 (10 ng/ml), BMP10 (10 ng/ml) or GI254023X (10 μM) for 24h. HUVECs were stained for VE-Cadherin (red), β-Catenin (green) using specific antibodies. Phalloidin was used to stain F-actin (blue). Scale bar = 30 μm

