## Supplementary figures and images for "Oxidative stress-induced MMP- and γ-secretase-dependent VE-cadherin processing is modulated by the proteasome and BMP9/10"

### Supplementary Fig.1

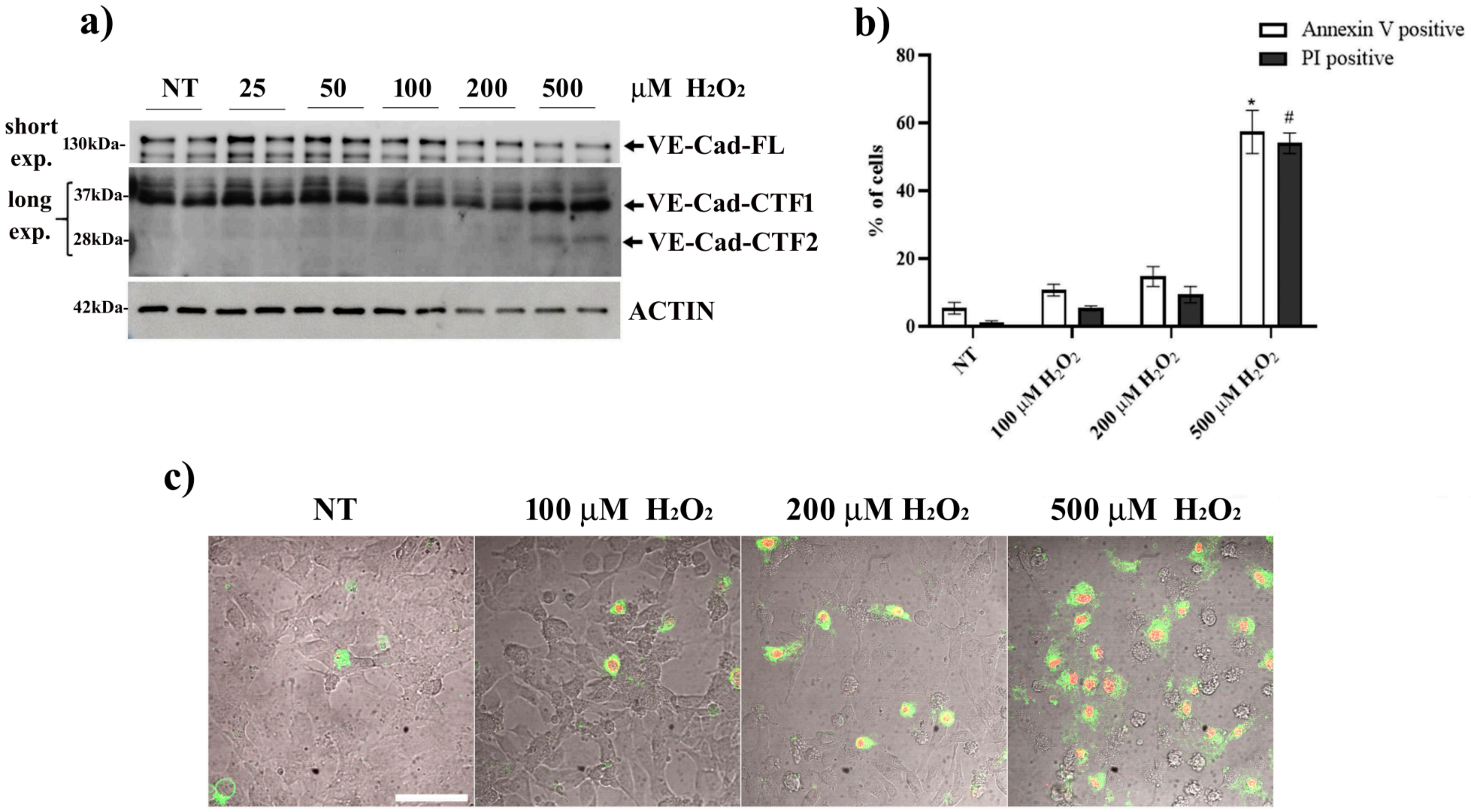


#### Supplementary Fig.1

### Supplementary Fig.2

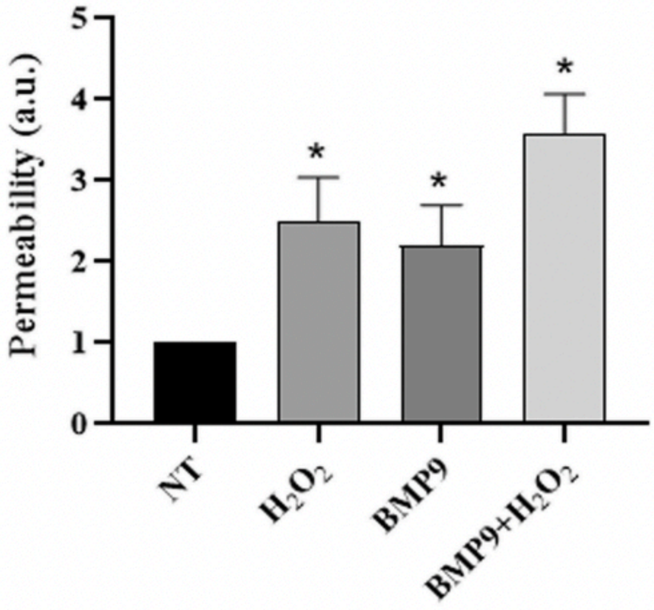


**Supplementary Fig.2**

### Supplementary Fig.3

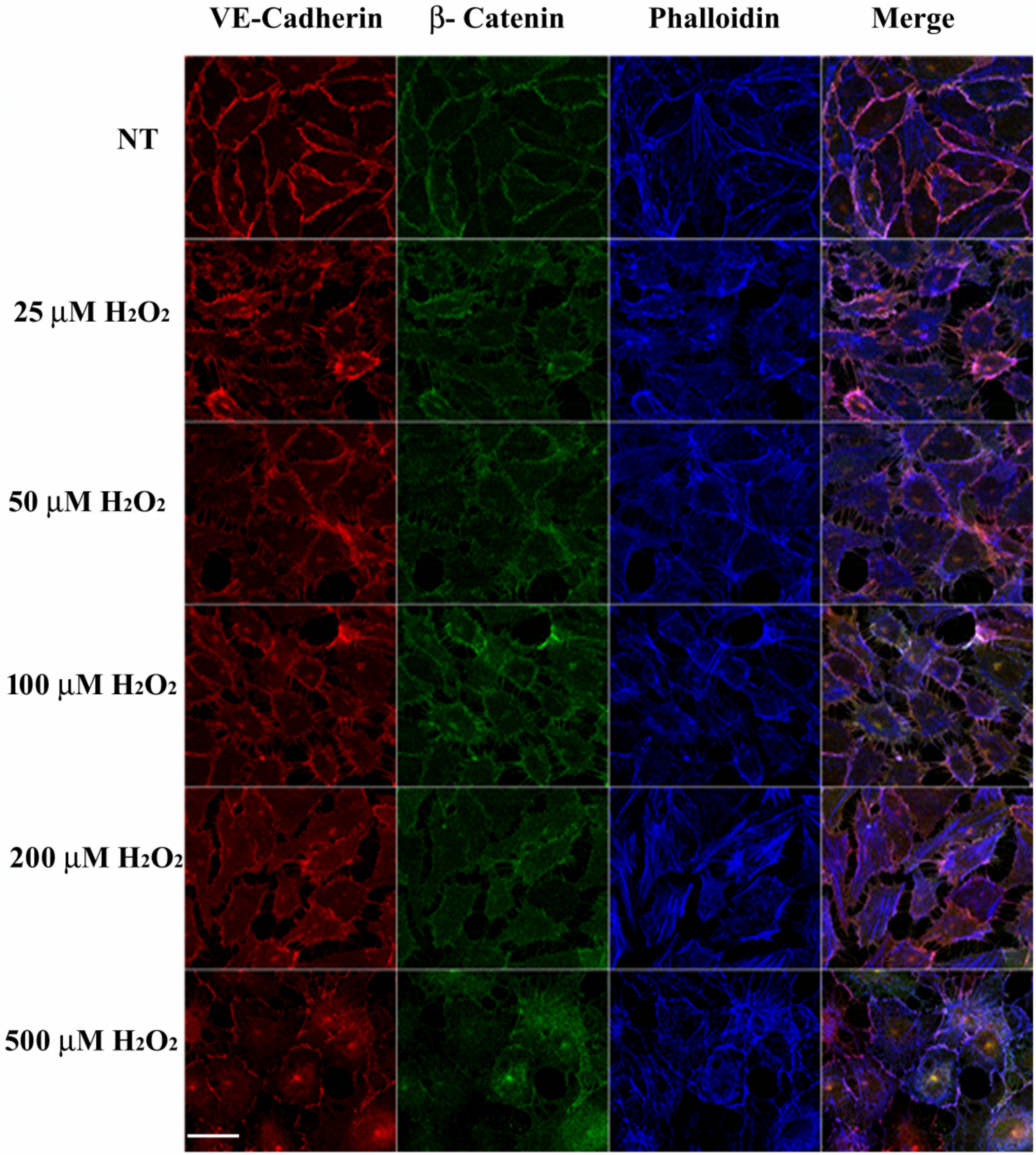


**Supplementary Fig.3**

### Supplementary Fig.4

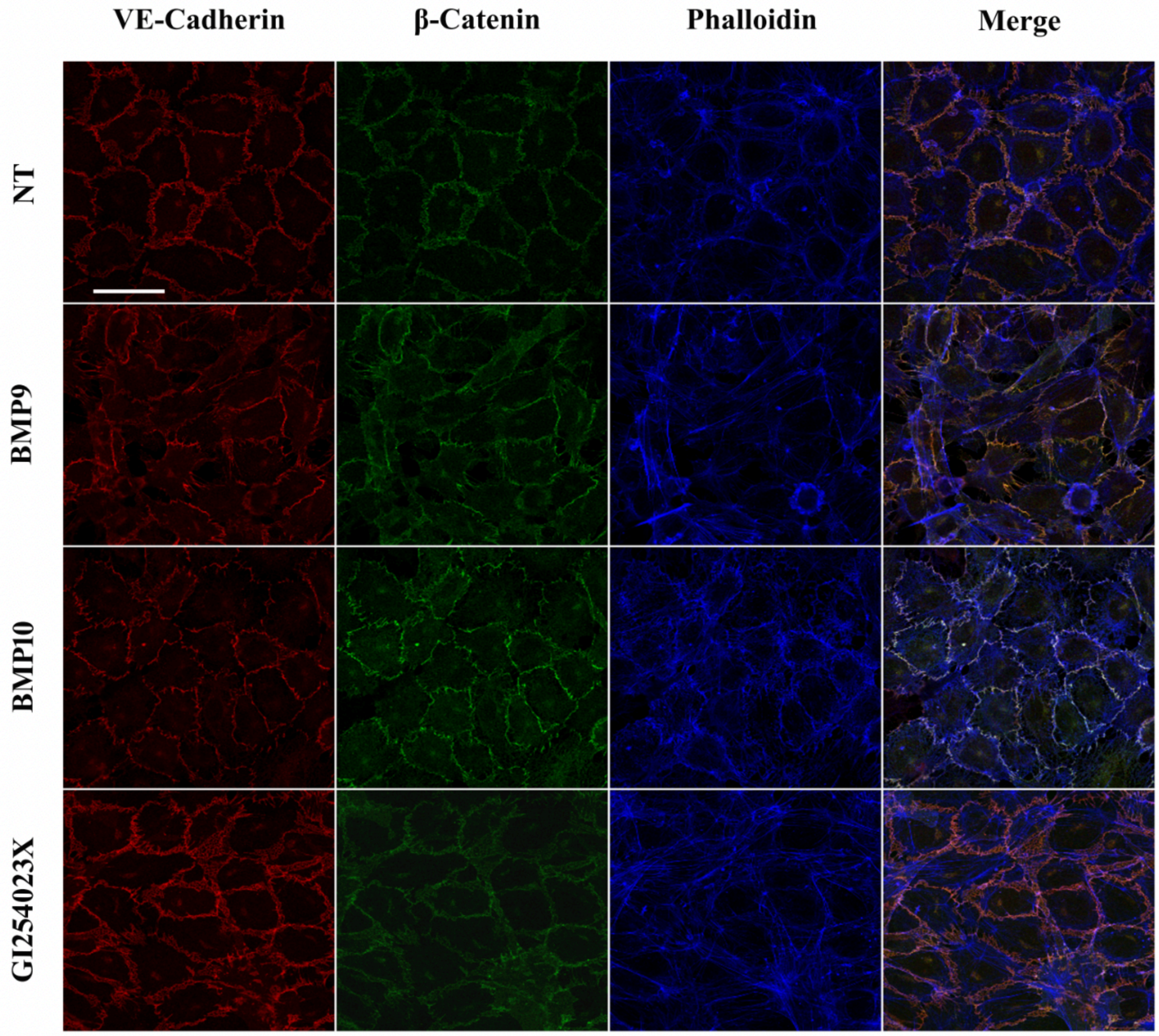


Supplementary Fig.4
